# Comparison of Small Gut and Whole Gut Microbiota of First-Degree Relatives with Adult Celiac Disease Patients and Controls

**DOI:** 10.1101/227272

**Authors:** Rahul Bodkhe, Sudarshan A. Shetty, Dhiraj P. Dhotre, Anil K. Verma, Khushbo Bhatia, Asha Mishra, Gurvinder Kaur, Pranav Pande, Dhinoth K. Bangarusamy, Beena P. Santosh, Rajadurai C. Perumal, Vineet Ahuja, Yogesh S. Shouche, Govind K. Makharia

## Abstract

Recent studies on celiac disease (CeD) have shown the role of gut microbiota alterations in CeD pathogenesis. Whether this alteration in the microbial community is the cause or effect of the disease is not well understood, especially in adult onset of disease. The first-degree relatives (FDRs) of CeD patients may provide an opportunity to study gut microbiome in pre-disease state as FDRs are genetically susceptible to CeD. By using 16S rRNA gene sequencing, we observed between the disease condition (CeD), pre-disease (FDR) and control subjects. However, differences were observed at the level of amplicon sequence variant (ASV), suggesting alterations in specific taxa between pre-diseases and diseased condition. Duodenal biopsies showed higher differences in ASVs compared to faecal samples indicating larger disruption of microbiota at disease site. Increased abundance of specific *Helicobacter* ASVs were observed in duodenum of CeD when compared to FDR (*p* < 0.01). In case of fecal samples CeD microbiome and *Actinomyces*. In addition, predicted functional metagenome showed reduced ability of gluten that ecosystem level diversity measures (except in the duodenum) were not significantly different is characterized by reduced abundance of beneficial taxa such as *Akkermansia, Ruminococcus* degradation by CeD faecal microbiota in comparison to FDRs and controls.

## Introduction

Celiac disease (CeD) is a common, chronic immune mediated enteropathy of the small intestine which affects approximately 0.7% of the global population (Singh et al., in press). Once thought to be uncommon in Asia, CeD is now prevalent in many Asian countries including India (Makharia et al., 2011). CeD is caused by the consumption of gluten proteins present in cereals such as wheat, barley and rye in genetically susceptible individuals (Caminero et al., 2015). While many genes are involved in the development of CeD, thus far only the presence of HLADQ2 or DQ8 haplotype is considered to be essential (Sanz and Pama, 2011). Additional factors that contribute to pathogenesis include other co-genetic factors (genome wide association studies have identified several markers), wheat-related factors (age of ingestion, type and quantity of wheat) and the way gluten is metabolized in the intestine (Kagnoff, 2007; van de Wal et al., 1998; Verdu et al., 2015). About 30-40% of the gluten protein consists of glutamine and proline. Since humans are unable to enzymatically break the molecular bonds between these two amino-acids, many immunogenic peptides are produced (Jabri and Sollid, 2006). There remains a possibility that enzymes secreted by the small intestinal microbiota convert some of these immunogenic peptides to non-immunogenic peptides.

While 20-30% of individuals in many countries including India are genetic susceptibility to develop CeD and the majority of them are exposed to wheat, only 1% of them develop CeD. This brings forth the role of other factors such as the gut microbiota in the pathogenesis of CeD (Sánchez et al., 2012). Recently, numerous studies have highlighted the potential role of gut microbiota in inflammatory gastrointestinal diseases (de Sousa Moraes et al., 2014; Fernandez-Feo et al., 2013; Png et al., 2010; Rivière et al., 2016; Schneeberger et al., 2015; Zeng et al., 2017).

However, these changes in the microbial community structure and function in patients with CeD are cause or effect of the disease state remains unclear to date. In order to answer this question, one has to examine the status of the gut microbiota in the pre-disease state. Recently two studies investigated the microbiota of at risk children who developed CeD few years after birth. One study observed an increase in *Bifidobacterium breve* and *Enterococcus* spp. in infants that developed active CeD (Olivares et al., 2018). Another study, did not observe any association between microbiota composition and development of CeD during the age of 9 and 12 months (Rintala et al., 2018). However, potential microbiota related triggers for development of CeD in later adult life still remain unclear. While 70-80% percent of first-degree relatives (FDRs) of patients with CeD have HLADQ2/DQ8 haplotype (compared to 30% in the general population); only approximately 8.5% of FDRs develop CeD (Singh et al., 2015). Thus, the question arises; why do only few FDRs develop CeD and what is the role of the gut microbiome in disease protection? Indirect evidence of altered microbiota in relatives of patients with CeD is suggested by significantly lower levels of acetic acid and total short chain fatty acids, and higher faecal tryptic activity (Tjellström et al., 2007). Nevertheless, to date there is no information on the gut microbial composition and function in FDRs of patients with CeD, especially using the latest sequencing approaches. Additionally, it is important to explore the status of the microbiota in both the small intestine, the site of the disease, and feces, as representative of whole gut microbiome. To test the hypothesis that gut microbiome of FDR is different from CeD and could potentially play an important role in the pathogenesis of CeD, we explored the composition of both small intestinal and the whole gut microbiome using Illumina MiSeq in a subset of patients with CeD, first degree relatives and controls. We further investigated the potential microbial functions that are characteristic of FDR and CeD microbiota.

## Results

### Comparison of faecal and duodenal microbial community in the study cohort

The characteristics of the study subjects have been summarized in the Table 1. All the participants were on staple gluten containing diet during sampling for this particular study. After diagnosis of CeD the patients underwent therapy with dietary recommendation to avoid gluten in daily diet. However, in the present study, we do not include samples after dietary changes. Both duodenal biopsies and faecal samples were included to investigate differences in both site-specific and whole gut bacterial diversity and community structure in patients with CeD, FDRs and controls. The microbial community was different between the faecal and duodenal biopsies irrespective of whether they were from CeD, FDR or DC groups (Supplementary figure S1a), (Analysis of similarities; ANOSIM statistic R: 0.4998, Significance: 0.001). Analysis of alpha diversity between the sampling sites suggested no significant differences between the sampling sites (Supplementary figure S1b). Further analyses were carried out separately for faecal and duodenal samples in different groups.

**Table 1:**
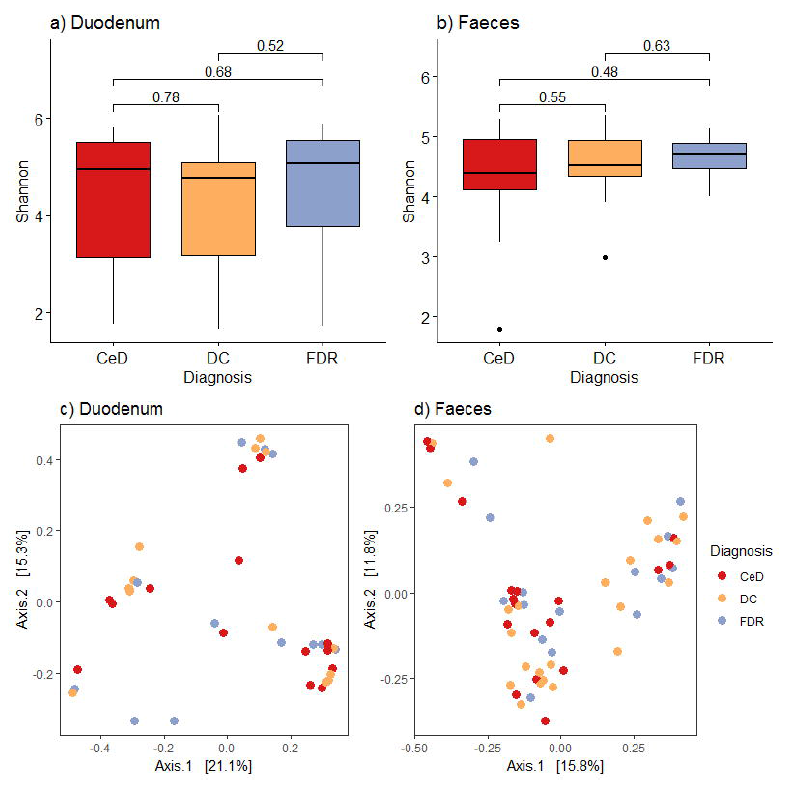
Demographic characteristics on study subjects

### Site specific bacterial community structure in FDRs, CeD and controls

#### Duodenal and faecal microbiota composition and structure is distinct in FDRs, CeD and control groups

To investigate if patients with CeD, FDRs or DC had site specific dissimilarities in microbiota composition, we analyzed microbiome composition of duodenal and faecal samples separately. Alpha diversity was determined using Shannon index, pairwise comparisons of alpha diversity in duodenal biopsies between FDRs, CeD and controls suggested no significant differences (Figure 1a). Similarly, for faecal samples no significant differences were observed for alpha diversity between diagnosis groups (Figure 1b).

**Figure 1:**
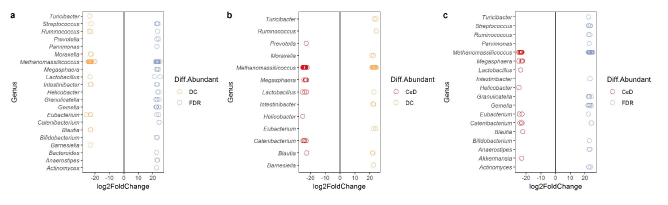
a. Comparison of alpha diversity between diagnosis groups in duodenal biopsies. b. Comparison of alpha diversity between diagnosis groups in Faecal samples. c. Principle coordinates analysis of bacterial community based on bray-curtis distance between diagnosis groups in duodenal biopsy samples. d. Principle coordinates analysis of bacterial community based on bray-curtis distance between diagnosis groups in faecal samples

Further, unconstrained comparison based on Bray-Curtis revealed no significant separation for duodenal biopsy microbiota between CeD, FDRs and control samples (Analysis of similarities; Anosim test; R-statistic =0.0014, p = 0.427 Figure 1c). In case of faecal microbiome, comparison based on Bray-Curtis distances between diagnosis groups was done. Similar to the duodenal biopsy microbiome it was not significantly different between diagnosis groups (Analysis of similarities; Anosim test; R-statistic = 0.051, p = 0.058 Figure 1d).

### Taxonomic differences in microbiota from duodenal biopsies of FDRs CeD, and controls

At phylum level in duodenal biopsy samples Actinobacteria, Bacteroides, Euryarchaeota, Firmicutes and Proteobacteria were the dominant members (Additional file figure S2). When performed pairwise comparison Actinobacteria (p= 0.013) and Bacteroides (0.02) were found be significantly increased in predisease state (FDR) in comparison to controls. Moreover, at order level FDR showed significant more abundance of Actinomycetales and Clostridiales than the control duodenal biopsies (p< 0.05) (Additional file figure S3).

To further investigate differences at lower taxonomic level between diagnosis groups, we used the DESeq2 package with default parameters.

### Changes in taxonomic abundance in the biopsies of FDRs in comparison to controls

Differential abundance analysis identified bacterial genera *Ruminococcus, Intestinibacter, Eubacterium* and *Anaerostipes* belonging to Clostridiales to be at least 21 fold higher in abundance in FDR biopsies (Figure 2a). Order Actinomycetales (p=0.02) and its genus *Actinomyces* were also observed in higher abundance in FDRs in comparison with controls. Notably, we observed differentially higher abundance of opportunistic pathogenic genera *Helicobacter* and *Prevotella* in duodenum of FDRs (>23 fold change, p < 0.01). In total a group 17 genera were significantly more abundant in FDR biopsy samples in comparison to control biopsies (p< 0.01) and these genera were at least 21 fold more in abundance. However, on the other side this analysis also identified 10 genera that were significantly depleted in FDR samples (>log2 Fold Change of 20, p < 0.01), including ASVs belonging to *Ruminococcus, Blautia, Eubacterium* and *Intestinibacter*. Among these the most significant difference in a bacterial genera was *Eubacterium* which was 26 fold decreased in FDR samples (p < 0.01).

**Figure 2:**
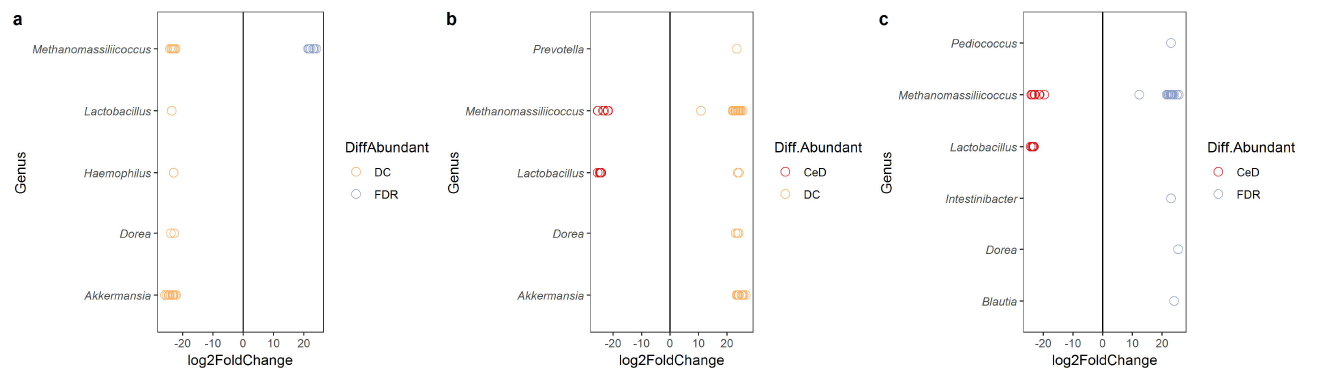
Comparison of differential abundance of bacterial taxa between the diagnosis groups in biopsy samples. a. Differential abundance DC vs FDR b. Differential abundance CeD vs DC c. Differential abundance CeD vs FDR. Only taxa with significant differences (P < 0.01) in log2 fold change are depicted.

### Changes in taxonomic abundance in the biopsies of patient with Celiac disease in comparison to controls

Next we compared microbial composition between CeD and controls to explore differentially abundant and reduced taxa in disease state. 35 ASVs were found to be at least 22-fold higher in abundance in duodenal biopsies of CeD group than the control biopsy samples. These ASVs were belonging to *Blautia, Catenibacter, Helicobacter, Lactobacillus, Megasphaera, Methanomassillicoccus* and *Prevotella* (Figure 2b). The most significant difference in a bacterial species were associated with *Lactobacillus, Methanomassiliicoccus, Catenibacter* and opportunistic pathogen *Helicobacter,* which were more than 22 fold higher in abundance in CeD biopsy samples than those of control samples (p < 0.01). Furthermore, *Megasphaera* and *Blautia* genera were also in higher abundance in CeD samples. Analysis also identified 34 ASVs belonging to 9 genera that were significantly depleted in CeD samples (p < 0.01). The majority of these genera (4/9) were belonging to the orders Clostridiales including *genera Ruminococcus, Intestinibacter, Blautia* and *Eubacterium*. Among these, the most depleted taxon was the short chain fatty acid (SCFA) producer *Ruminococcus*, which was 24 fold reduced in samples from those with CeD (p< 0.01). Moreover, higher abundance of genus *Turicibacter,* and *Moraxella* was significantly associated with a control microbial configuration in comparison with CeD.

### Changes in taxonomic abundance in the biopsies of patient with Celiac disease in comparison to First degree relatives of CeD

Next, to identify the differentially abundant taxa between predisease and disease state we did similar analysis for CeD and FDR groups. DESeq2 identified a group of 27 taxa belonging to Firmicutes and Proteobacteria that were significantly more abundant in CeD duodenal samples. These taxa were found to be at least 22-fold higher in abundance and were belonging to genera *Blautia, Eubacterium, Helicobacter, Lactobacillus, Megasphaera* and *Akkermansia*. Similar to the comparison with controls, bacterial genera *Methanomassillicoccus, Catenibacter* and *Helicobacter* were the most significantly abundant bacterial genera in CeD duodenum samples in comparison to FDRs (>24 fold change, p < 0.01). In addition, *Moraxella* and *Eubacterium* were also the other most differential abundant taxa were associated with duodenum in disease condition (>24 fold change, p < 0.01).

We also identified 59 taxa belonging Firmicutes and Actinobacteria were significantly depleted in CeD samples (p < 0.01). Also the order Clostridiales and the beneficial genera affiliated to it such as *Ruminococcus, Intestinibacter* and *Anaerostipes* were also significantly reduced in CeD biopsies.

Moreover, *Gemella* a commensal genus of the upper respiratory tract, gluten degrader *Actinomyces* and genera *Streptococcus* and *Bifidobacterium* were also found to be significantly low in abundance in CeD (Figure 2c).

### Taxonomic differences in the faecal microbiota in patients with CeD, FDRs and controls

Phylum level comparison of microbial community between CeD, FDRs and controls demonstrated that Proteobacteria, Actinobacteria, Bacteroidetes, Euryarchaeota and Firmicutes constitute the majority of the faecal microbiota (Additional file figure S4). However in contrast to the biopsy samples Bacteroidetes was found to be marginally decreased in FDR samples in comparison to controls (p= 0.058). Similar trend was observed for order Bacteroidales, it showed marginal lower abundance in FDRs (p= 0.054). However, order Clostridiales was significantly abundant in FDRs in comparison to controls (p= 0.017) (Additional file figure S5).

### Changes in taxonomic abundance in the feces of FDR in comparison with Controls

In faeces of FDRs mostly the significant depletion (22/30) of beneficial taxa was observed. Only the archaeal genus *Methanomassiliicoccus* was observed differentially abundant in FDRs faecal samples than those of control samples (Figure 3a). However, 7 ASVs belonging to same genus were significantly reduced in FDRs. Further analysis identified more than 23 fold (p < 0.01) reduction in bacterial genera which are known for a healthy microbiota homeostasis, which include *Akkermansia, Lactobacillus* and *Dorea*.

**Figure 3:**
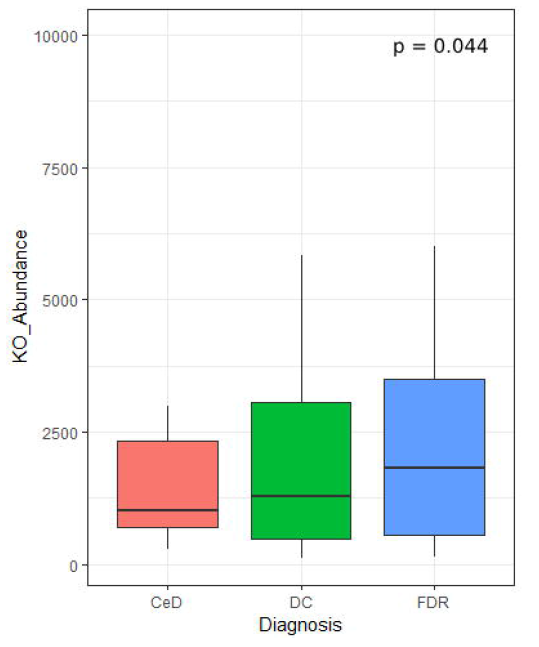
Comparison of differential abundance of bacterial taxa between the diagnosis groups in faecal samples. a. Differential abundance CeD vs DC. b. Differential abundance CeD vs FDR. Only taxa with significant differences (P < 0.01) in average log2 fold change are depicted.

### Changes in taxonomic abundance in the feces of CeD in comparison with Controls

Similar to the FDRs, mostly the depletion of bacterial taxa was observed in CeD faecal samples when compared with controls. Moreover, the same ASVs of *Akkermansia, Lactobacillus* and *Dorea* were significantly depleted in CeD (Figure 3b). In addition *Prevotella* showed 23 fold (p< 0.01) reductions in abundance in CeD. On the other hand, DESeq2 identified genus *Lactobacillus* to be in significant abundance in disease condition (CeD) in comparison to control fecal samples.

### Changes in taxonomic abundance in the feces of CeD in comparison with FDRs

To explore differentially abundant taxa in disease condition in comparison to predisease state, we compared microbial composition between CeD and FDRs. In disease state mostly a significant depletion was observed for physiologically important bacterial taxa compared with FDRs faeces (Figure 3c). Order Clostridiales and genera *Intestinibacter, Dorea* and *Blautia* belonging to this order were significantly in lower abundance in CeD. In addition, *Pediococcus* was found to be 23 fold reduced abundance in CeD, however ASVs affiliated with *Lactobacillus* were more than 24 fold differentially abundant in CeD in comparison FDRs.

### Imputed metagenome of FDR and CeD duodenal microbiome shows reduced proportion of genes involved in gluten metabolism in comparison to that of the controls

In addition to differentially abundant microbial taxa, different study groups might have altered metabolic potential. Of specific interest were the enzymes related to glutenases as they play a role in breakdown of gliadin residues. We followed Piphillin workflow to predict functional profile of fecal microbial community (Iwai et al., 2016). A total of 159 KEGG orthologies (KO) were significantly different between diagnosis groups (Supplementary Table 1). Among these the KO abundance for Xaa-pro dipeptidase (K01271, Prolidase) enzyme which is known to have role in gluten degradation was found to be significantly reduced in CeD as compared to FDR and controls (figure 4).

**Figure 4:**
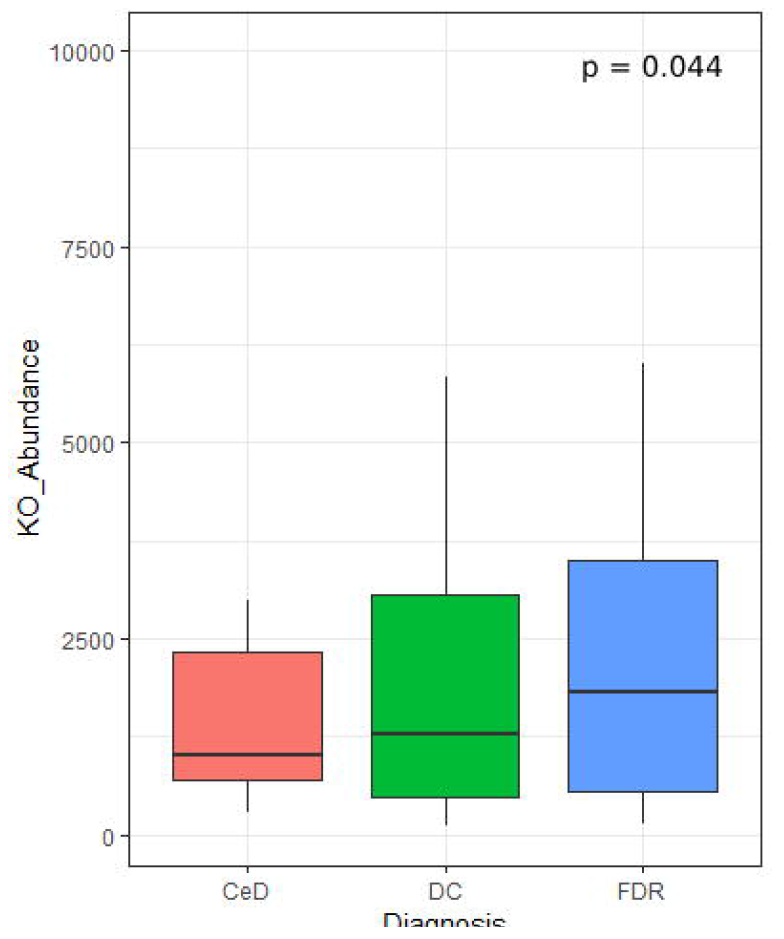
KO abundance for Xaa-pro dipeptidase (K01271) enzyme in faeces inferred from predicted metagenome for faecal samples. Comparison was done using ANOVA.

## Discussion

The aim of the present study was to investigate differences in the duodenal and faecal microbiota of pre-diseased state i.e. FDRs subjects compared to diseased state i.e. CeD and controls. The FDR group was included for two main reasons: 1) They represent a population which is genetically-susceptible to develop CeD; 2) They provide a unique opportunity to identify features of the host as well as of the associated microbiota that may be involved in the protection against developing CeD. We collected both duodenal biopsies and faecal samples to investigate both local and overall changes in the microbiota in FDR, patients with CeD and controls.

To the best of our knowledge, reports on site specific microbiota patterns in adult patients with CeD remain scarce, and no results on both site specific and whole gut microbiome on FDRs have been reported to date. Present study provides an overall view on differences of both site-specific changes as well as changes in the faecal microbiota of FDRs, CeD and DC.

At lower taxonomic level, several taxa were identified to be differentially abundant between the diagnosis groups. Notably, ASVs related to *Helicobacter, Ruminococcus*, *Megasphaera,* and *Lactobacillus,* showed higher (> 24) log2 fold change in CeD biopsy samples. When we performed analysis at species level, abundance of an ASV1811, *H. pylori* was higher in CeD compared to controls and FDR subjects. In turn FDR were found to harbor higher abundance of ASV2016 and ASV4095 belonging to *H. pylori* in comparison to controls (Supplementary information). As per previous reports, CeD patients with *H. pylori* gastritis were found with increased numbers of intraepithelial lymphocytes in the duodenal mucosa (Villanacci et al., 2006). In contrast, there are also reports which failed to reveal a relationship between *H. pylori* and CeD and found that *H. pylori* presence is inversely associated CeD (Lebwohl et al., 2013). However, in our study the ASV which is abundant in CeD is different than those ASVs which are enriched in FDR. Our analysis included finer sequence level variation and differentiated single nucleotide level difference (Callahan et al., 2016). In view of intra-genomic differences in 16S rRNA gene, we compared the 16S rRNA gene copies of *H. pylori* in publically available genomes. We observed that on average the *H. pylori* genome has two copies of 16S rRNA gene and we did not observe differences between the two copies within a single genome in the V4 region investigated here (Supplementary information text page 3-7). Therefore, future studies need to focus on strain level variations and functional aspects of *H. pylori* in regards with CeD using metagenomics and functional omics. Other ASVs which showed higher abundance in CeD as compared to FDR and DC and lower abundance in FDR as compared to DC are *Megasphaera* and *Catenibacterium*. One more important observation from differential abundance results is decreased abundance of *Ruminococcus* in CeD as compared to FDR and CeD. *Ruminococcus* is considered to be a keystone taxa with influence on the microbial community and responsible for the major fraction of butyrate production in the gut (Morrison and Preston, 2016; Shetty et al., 2017; Ze et al., 2012).

Duodenal microbiota of FDR is characterized by increased abundance of ASVs related to *Actinomyces, Streptococcus, Bifidobacterium* and *Anaerostipes*. These taxa are known to possess gluten degrading enzymes, probiotic properties and ability to produce SCFA respectively (Barrangou et al., 2009; Couvigny et al., 2015; Fernandez-Feo et al., 2013; Morrison and Preston, 2016; Rivière et al., 2016). Moreover, the strain belonging to *Bifidobacterium* was reported to prevent gluten-related immunopathology in mice (McCarville et al., 2017). Higher number of these taxa in small intestine of FDRs may indicate protective role of these taxa in pre-disease state.

In comparison to duodenal biopsies, less numbers of ASVs were differentially abundant between diagnosis groups in faecal samples. This indicates more disrupted microbiome at disease site than overall gut microbiome and highlights the importance of inclusion of biopsy samples in present study.

In faecal samples, CeD showed significant enrichment of ASVs affiliated to genus *Lactobacillus* in comparison to FDR and DC. This is in line with previous study in which, higher abundance of *Lactobacillus* was observed in oral microbiome of patients with CeD (Tian et al., 2017). Moreover, there are reports stating that the certain *Lactobacillus* species degrade gliadin and increases the availability of antigenic peptides (Engstrom et al., 2015).In the present study, higher abundance of *Lactobacillus* in CeD microbiota may indicate their ability to breakdown gluten into pro-inflammatory peptides in the small intestine. Another important observation from differential abundance is the lower abundance of *Dorea* and *Akkermansia* in faecal sample of CeD. Both of these taxa are known to produce SCFAs which in turn strengthens the health of enterocytes and inhibits intestinal inflammation (Ohira et al., 2017).

Lower abundance of *Lactobacillus* and higher abundance of *Dorea* in FDR and DC in comparison to CeD indicates that the faecal microbiota of FDR is more similar to the microbiota of control samples. However, *Akkermansia* ASVs were in more abundance in control as compared to FDR.

Overall, microbiota of DC can be characterized with enriched abundance of SCFA producing core bacterial taxa, such as *Ruminococcus* and *Akkermansia* in case of biopsy and faecal samples respectively. ASVs such as *Megasphaera, Ruminococcus* and *Helicobacter* in duodenal biopsy of FDR showed higher abundance when compared with control but they are reduced in comparison with disease state CeD. Moreover, *Dorea* showed similar abundance pattern in faecal samples. In our study, the higher abundance of known pathogenic bacteria such as *Helicobacter* (well-known bacteria to be associated with intestinal inflammation) and reduced abundance of health associated bacteria such as *Akkermansia, Ruminococcus* (bacteria known for the anti-inflammatory properties) and *Actinomyces* (a well-known gluten degrader) emerged as the characteristic of the CeD microbiota.

Through metagenome prediction method we found that the gene abundance for Xaa-pro Dipeptidase enzymes was less in CeD as compared to FDR and controls. This enzyme shows a high specificity for proline residues present in gluten and hydrolyze the peptide bond (Park et al., 2004). These observations suggest that the FDR and CeD duodenal microbiota differs in the bacterial composition and that loss or gain of specific bacteria capable of gluten degradation. This may impact gluten processing and the presentation of immunogenic gluten epitopes to the immune system in the small intestine. However, link between the predicted metagenome and gut microbiome needs to be validated with *in-vitro* enzyme assay.

The present study was conducted to investigate if the duodenal and faecal microbiotas of FDR of CeD patients are different from that of CeD and controls. Our observations from PCoA (Figure 1c and 1d) and differential abundance (Figure 2 and 3) suggest variation at lower taxonomic levels. These potential species and/or strain level variations and functional aspects need to be investigated using shotgun metagenomics and functional omics in follow-up studies.

However, metagenomic studies of biopsy samples remain a challenge because of high proportion of host DNA. Thus, predictive metagenomics using 16S rRNA gene as a practical solution was employed for biopsies. In this initial exploratory study, we investigated the gut microbiome with respect to the disease status only and future studies considering other confounding factors such as diet, body mass index age, sex, frequency and quantity of gluten intake among others will be required for a better understanding the gut microbiome in CeD and FDRs. Additionally, the control group in our study was not healthy subjects but patients with functional dyspepsia. These subjects were used as proxy since invasive sampling procedures such as endoscopy from clinically healthy subjects is not permitted under the institutional regulations.

In summary, present study highlights the specific differences in the microbiota of FDR compared to that in patients with CeD and controls. Difference in FDR microbiota in both the faecal and duodenal biopsy samples compared to CeD suggests microbiota of FDR have unique features. Analysis of single nucleotide level variation provides a finer resolution and suggests that changes in strain level features need to be investigated in CeD. These unique features should be addressed in future mechanistic studies to understand etiopathogenesis of CeD.

## Conclusions

Significant differences at ASV level suggest that specific bacterial taxa like *Helicobacter* may be important for pathogenesis of CeD. Higher abundance of beneficial bacterial taxa especially SCFA producers in controls suggest that there may be a protective role of these taxa in CeD development. Moreover, the predicted differences in gluten metabolism potential by FDR and CeD microbiota point towards the need for investigating functional capabilities of specific bacterial taxa in healthy FDR and CeD patients.

## Methods

### Patients and Methods

#### Human subjects, duodenal biopsies and faecal sample collection

A total of 62 subjects participated in this study including 23 treatment naïve patients with CeD [all HLA-DQ2/DQ8+, having high titre of anti-tissue transglutaminase antibodies (tTG Ab) and having villous abnormalities of modified Marsh grade 2 or more], 15 healthy first-degree relatives of patients with CeD [having normal titre of anti-tTG Ab and having no villous abnormalities of modified Marsh grade 0 or 1], and 24 controls (patients with Hepatitis B Virus carriers or those having functional dyspepsia; having normal titre of anti-tTG Ab and having no villous abnormalities) (Table 1). Duodenal biopsies and faecal samples were collected from each of the above mentioned subjects at All India Institute of Medical Sciences, New Delhi, and sent to National Centre for Cell Sciences, Pune for microbiome analysis. The ethics committees of All India Institute of Medical Sciences, New Delhi, and National Centre for Cell Sciences, Pune, India approved the study. Informed and written consent was obtained from all the participants. Further details of patients and controls have been provided in the (Supplementary file 1: Table 2).

#### DNA extraction and 16S rRNA gene sequencing

Total DNA was extracted from duodenal biopsies using QIAGEN DNeasy Blood and Tissue kit (QIAGEN, Germany) and faecal samples using the QIAamp fast DNA stool Mini Kit (QIAGEN, Germany) according to the manufacturer’s instructions. We used Illumina MiSeq sequencing to determine the bacterial composition of the duodenal biopsies and faecal samples. PCR was set up in 50 μl reaction using AmpliTaq Gold PCR Master Mix (Life Technologies, USA) and with 16S rRNA V4 variable region specific bacterial primers 515F (5′-GTGCCAGCMGCCGCGGTAA-3′) and 806R (5′‐ GGACTACHVGGGTWTCTAAT-3′)

#### Sequence processing and bacterial community analysis

Illumina Miseq platform rendered a total of 76058052 raw 16S rRNA sequence reads for the 102 faecal and biopsy samples of the diagnosis groups, with an average of 745667 ±194667 reads per sample. Adapter sequences were trimmed by using Cutadapt (1.18) tool (Martin, 2011)and trimmed reads were DADA2 (v 1.6.0) pipeline (Callahan et al., 2016). In the first step reads were inspected for read quality profile, the read quality score was decreased (<30) after 240 bases for forward read and 160 bases for reverse reads. We truncated the forward reads at position 240 (trimming the last 10 nucleotides) pooled as Fasta.gz file format for further analysis in and reverse reads at position 160 (trimming the last 90 nucleotide). After quality filtering and removal of bases with a total of 70502947 (92.69%) high-quality reads of the 16S rRNA amplicons were obtained, with an average 691205 ± 181263 reads per sample, ranging from 325350 to 1207169 among samples (Supplementary Table 3). Finally, taxonomic assignment was done by the naive Bayesian classifier method with default setting as implemented in DADA2, against Human Intestinal 16S rRNA gene reference taxonomy database (HITdb v 1.00). Briefly, HITdb is a 16S rRNA gene database based on high-quality sequences specific for human intestinal microbiota, this database provides improved taxonomic up to the species level (Ritari et al., 2015). Unassigned chimeric and sequences of chloroplast and mitochondria were excluded from downstream analysis. Taxonomic assignment successfully mapped 6567144 ASVs (Amplicon Sequence Variants), with an average of 64383 ± 29929 ASVs per sample. Finally, from these ASVs, ASV table was constructed and the ASVs generated by the contaminants were removed by using decontam software (Davis et al., 2017) and the output ASV table was used for downstream analyses.

Microbial diversity and composition analysis was done using the R-package phyloseq (v1.22.3) (McMurdie and Holmes, 2013) and microbiome R package (v1.0.2) (Leo and Shetty, 2017). To test for similarities in bacterial communities between sample types and diagnosis groups Analysis of Similarities (ANOSIM) on Bray-Curtis distances was used. ANOSIM is a function in vegan package (v 2.4-4) to calculate significance of PCoA clustering based on the Bray-Curtis distances (Dixon, 2003).

To identify differentially abundant ASVs in pairwise comparisons between diagnosis groups we used DESeq2 (v1.18.0) (Love et al., 2014). All ASVs that were significantly (alpha = 0.01) different in abundance between the diagnosis groups were reported and were adjusted for multiple comparisons using the Benjamini-Hochberg, false discovery rate procedure. Data was visualized using ggplot2 (v 2.2.1) in R (Hadley Wickham, 2016).

### Metagenomic Imputation

Piphillin tool was used to infer metagenome from 16S rRNA ASV counts table and representative sequence of each ASV. Briefly, this tool predicts metagenomes with high accuracy by leveraging the most-current genome reference databases (Iwai et al., 2016). It uses direct nearest-neighbor matching between 16S rRNA amplicons and genomes to predict the represented genomes. Latest version (May 2017) of KEGG database and 97% of the identity cutoff was selected for the prediction. The output from Piphillin was further analyzed by STAMP statistical tool, ANOVA with post hoc Tukey-kramer test was used to identify statistically different KEGG orthologies between diagnosis groups (Parks et al., 2014).

## List of abbreviations

CeD: Celiac disease
DC: Diseased controls (dyspeptic)
FDR: First degree relatives
ASV: Operational taxonomic unit
rRNA: Ribosomal Ribonucleic acid
PCoA: Principal coordinates analysis

## Declarations

## Acknowledgement

We acknowledge the help of Aishwairya Sharma, a dietician, who counselled the patients for gluten free diet. The authors are grateful to Dr. Nachiket Marathe for fruitful discussion related to the project. The authors thank Johanna Gutleben for proofreading the manuscript. We sincerely acknowledge the support of each participant of this study who accepted to participate in this study.

## Availability of data and materials

Sequence data generated in this study is available from the NCBI Sequence Read Archive within the Bioproject ID accession PRJNA385740. (https://www.ncbi.nlm.nih.gov/bioproject/?term=PRJNA385740) and to reproduce the analysis done in R, the R Markdown file and required data are available at https://github.com/rahulnccs/Comparison-of-Small-Gut-and-Whole-Gut-Microbiota-of-First-Degree-Relatives-with-Adult-Celiac-Disease.

Conflicts of interest: All the authors disclose no conflict of interest

## Authors Contributions

The research study was conceptualized, designed and supervised by GKM, YSS and VA. Patient recruitment, diagnosis and endoscopic examination was done by GKM; HLA testing was done by GK; biological sample collection (duodenal biopsy/stool) storage and maintenance was done by AKV, KB and AM. The extraction of genomic DNA was done by RB and PP. DKB, BPS and RCP were involved in amplicon sequencing. Bioinformatics analysis for amplicon data was done by SAS, DPD and RB. Data acquisition, data interpretation and drafting of the manuscript was done by SAS and GKM. YSS, DPD and VA critically reviewed the manuscript. All authors have read and approved the final manuscript.

## Ethics approval and consent to participate

The Ethics Committees of All India Institute of Medical Sciences, New Delhi, and National Centre for Cell Sciences, Pune, India approved the study. Informed and written consent was obtained from all the participants.

Consent for publication

Not applicable.

Supplementary file, Figure S1: a. Principal coordinates analysis (PCoA) of bacterial community in the faecal and duodenal biopsies based on Bray–Curtis distance. b. Comparison of alpha diversity measures between sampling sites.

Supplementary file, Figure S2: Phylum level distribution of ASVs in biopsy samples. Pairwise comparisons were done using Wilcoxon tests.

Supplementary file, Figure S3: Order level distribution of ASVs in biopsy samples. Pairwise comparisons were done using Wilcoxon tests.

Supplementary file, Figure S4: Phylum level distribution of ASVs in faecal samples. Pairwise comparisons were done using Wilcoxon tests.

Supplementary file, Figure S5: Order level distribution of ASVs in faecal samples. Pairwise comparisons were done using Wilcoxon tests.

Supplementary Table 1: Significantly different KEGG orthologies (KO) between diagnosis groups

Supplementary Table 2: Details of study subjects

Supplementary Table 3: Number of reads per sample at each stage of analysis. Supplementary Information document: Differential Abundance of Amplicon Sequence Variant of *Helicobacter.* Multiple sequence alignment was performed by CLUSTAL 2.0.11

## References

Barrangou R., Briczinski E. P., Traeger L. L., Loquasto J. R., Richards M., Horvath P., et al. (2009). Comparison of the Complete Genome Sequences of Bifidobacterium animalis subsp. *lactis DSM 10140 and Bl-04 †.* J. Bacteriol. 191, 4144–4151. doi: 10.1128/JB.00155-09.

Callahan B. J., McMurdie P. J., Rosen M. J., Han A. W., Johnson A. J. A., and Holmes S. P. (2016). DADA2: High-resolution sample inference from Illumina amplicon data. Nat. Methods 13, 581–583. doi: 10.1038/nmeth.3869.

Caminero A., Nistal E., Herrán A. R., Pérez-Andrés J., Ferrero M. A., Vaquero Ayala L., et al. (2015). Differences in gluten metabolism among healthy volunteers, coeliac disease patients and first-degree relatives. Br. J. Nutr. 114, 1157–1167. doi: 10.1017/S0007114515002767.

Couvigny B., de Wouters T., Kaci G., Jacouton E., Delorme C., Doré J., et al. (2015). Commensal Streptococcus salivarius Modulates PPARγ Transcriptional Activity in Human Intestinal Epithelial Cells. PLoS One 10, e0125371. doi: 10.1371/journal.pone.0125371.

Davis N. M., Proctor D., Holmes S. P., Relman D. A., and Callahan B. J. (2017). Simple statistical identification and removal of contaminant sequences in marker-gene and metagenomics data. bioRxiv, 221499. doi: 10.1101/221499.

de Sousa Moraes L. F., Grzeskowiak L. M., de Sales Teixeira T. F., and do Carmo Gouveia Peluzio M. (2014). Intestinal microbiota and probiotics in celiac disease. Clin. Microbiol. Rev. 27, 482–489. doi: 10.1128/CMR.00106-13.

Dixon P. (2003). VEGAN, a package of R functions for community ecology. J. Veg. Sci. 14, 927–930. doi: 10.1111/j.1654-1103.2003.tb02228.x.

Engstrom N., Sandberg A. S., and Scheers N. (2015). Sourdough fermentation of wheat flour does not prevent the interaction of transglutaminase 2 with??2-gliadin or gluten. Nutrients 7, 2134–2144. doi: 10.3390/nu7042134.

Fernandez-Feo M., Wei G., Blumenkranz G., Dewhirst F. E., Schuppan D., Oppenheim F. G., et al. (2013). The cultivable human oral gluten-degrading microbiome and its potential implications in coeliac disease and gluten sensitivity. Clin. Microbiol. Infect. 19, 1–14. doi: 10.1111/1469-0691.12249.

Hadley Wickham (2016). ggplot2: Elegant Graphics for Data Analysisn info. Springer-Verlag New York Available at: https://cran.r-project.org/web/packages/ggplot2/citation.html [Accessed July 31, 2018].

Iwai S., Weinmaier T., Schmidt B. L., Albertson D. G., Poloso N. J., Dabbagh K., et al. (2016). Piphillin: Improved Prediction of Metagenomic Content by Direct Inference from Human Microbiomes. PLoS One 11, e0166104. doi: 10.1371/journal.pone.0166104.

Jabri B., and Sollid L. M. (2006). Mechanisms of Disease: immunopathogenesis of celiac disease. Nat. Clin. Pract. Gastroenterol. Hepatol. 3, 516–525. doi: 10.1038/ncpgasthep0582.

Kagnoff M. F. (2007). Celiac disease: pathogenesis of a model immunogenetic disease. J. Clin. Invest. 117, 41–9. doi: 10.1172/JCI30253.

Lebwohl B., Blaser M. J., Ludvigsson J. F., Green P. H. R., Rundle A., Sonnenberg A., et al. (2013). Decreased risk of celiac disease in patients with Helicobacter pylori colonization. Am. J. Epidemiol. 178, 1721–30. doi: 10.1093/aje/kwt234.

Leo L., and Shetty S. (2017). microbiome R package. Bioconductor.

Love M. I., Huber W., and Anders S. (2014). Moderated estimation of fold change and dispersion for RNA-seq data with DESeq2. Genome Biol. 15, 550. doi: 10.1186/s13059-014-0550-8.

Makharia G. K., Verma A. K., Amarchand R., Bhatnagar S., Das P., Goswami A., et al. (2011). Prevalence of celiac disease in the northern part of India: A community based study. J. Gastroenterol. Hepatol. 26, 894–900. doi: 10.1111/j.1440-1746.2010.06606.x.

Martin M. (2011). Cutadapt removes adapter sequences from high-throughput sequencing reads. EMBnet.journal 17, 10. doi: 10.14806/ej.17.1.200.

McCarville J. L., Dong J., Caminero A., Bermudez-Brito M., Jury J., Murray J. A., et al. (2017). A commensal Bifidobacterium longum strain improves gluten-related immunopathology in mice through expression of a serine protease inhibitor. Appl. Environ. Microbiol. 83, e01323-17. doi: 10.1128/AEM.01323-17.

McMurdie P. J., and Holmes S. (2013). Phyloseq: An R Package for Reproducible Interactive Analysis and Graphics of Microbiome Census Data. PLoS One 8, e61217. doi: 10.1371/journal.pone.0061217.

Morrison D. J., and Preston T. (2016). Formation of short chain fatty acids by the gut microbiota and their impact on human metabolism. Gut Microbes 7, 189–200. doi: 10.1080/19490976.2015.1134082.

Ohira H., Tsutsui W., and Fujioka Y. (2017). Are Short Chain Fatty Acids in Gut Microbiota Defensive Players for Inflammation and Atherosclerosis? J. Atheroscler. Thromb. 24, 660–672. doi: 10.5551/jat.RV17006.

Olivares M., Walker A. W., Capilla A., Benítez-Páez A., Palau F., Parkhill J., et al. (2018). Gut microbiota trajectory in early life may predict development of celiac disease. Microbiome 6, 36. doi: 10.1186/s40168-018-0415-6.

Park M.-S., Hill C. M., Li Y., Hardy R. K., Khanna H., Khang Y.-H., et al. (2004). Catalytic properties of the PepQ prolidase from Escherichia coli. Arch. Biochem. Biophys. 429, 224–230. doi: 10.1016/J.ABB.2004.06.022.

Parks D. H., Tyson G. W., Hugenholtz P., and Beiko R. G. (2014). STAMP: statistical analysis of taxonomic and functional profiles. Bioinformatics 30, 3123–4. doi: 10.1093/bioinformatics/btu494.

Png C. W., Lind S. K., Gilshenan K. S., Zoetendal E. G., Mcsweeney C. S., Sly L. I., et al. (2010). Mucolytic Bacteria With Increased Prevalence in IBD Mucosa Augment In Vitro Utilization of Mucin by Other Bacteria. Am J Gastroenterol 2010; 105, 2420–2428. doi: 10.1038/ajg.2010.281.

Rintala A., Riikonen I., Toivonen A., Pietilä S., Munukka E., Pursiheimo J.-P., et al. (2018). Early fecal microbiota composition in children who later develop celiac disease and associated autoimmunity. Scand. J. Gastroenterol. 53, 403–409. doi: 10.1080/00365521.2018.1444788.

Ritari J., Salojärvi J., Lahti L., and de Vos W. M. (2015). Improved taxonomic assignment of human intestinal 16S rRNA sequences by a dedicated reference database. BMC Genomics 16, 1056. doi: 10.1186/s12864-015-2265-y.

Rivière A., Selak M., Lantin D., Leroy F., and De Vuyst L. (2016). Bifidobacteria and butyrate-producing colon bacteria: Importance and strategies for their stimulation in the human gut. Front. Microbiol. 7. doi: 10.3389/fmicb.2016.00979.

Sánchez E., Laparra J. M., and Sanz Y. (2012). Discerning the role of bacteroides fragilis in celiac disease pathogenesis. Appl. Environ. Microbiol. 78, 6507–6515. doi: 10.1128/AEM.00563-12.

Sanz Y., and Pama G. De (2011). Unraveling the Ties between Celiac Disease and Intestinal Microbiota. Int Rev Immunol 30, 207–218. doi: 10.3109/08830185.2011.599084.

Schneeberger M., Everard A., Gómez-Valadés A. G., Matamoros S., Ramírez S., Delzenne N. M., et al. (2015). Akkermansia muciniphila inversely correlates with the onset of inflammation, altered adipose tissue metabolism and metabolic disorders during obesity in mice. Sci. Rep. 5, 16643. doi:10.1038/srep16643.

Shetty S. A., Hugenholtz F., Lahti L., Smidt H., and de Vos W. M. (2017). Intestinal microbiome landscaping: insight in community assemblage and implications for microbial modulation strategies. FEMS Microbiol. Rev. 41, 182–199. doi:10.1093/femsre/fuw045.

Singh P, Arora A, Strand TA, Leffler D, Catassi C, Green PH, Kelly CP, Ahuja V M. G., and G Global prevalence of celiac disease: Systematic review and Meta-analysis. Gastroenterol Hepatol.(in press)

Singh P., Arora S., Lal S., Strand T. A., and Makharia G. K. (2015). Risk of Celiac Disease in the First‐ and Second-Degree Relatives of Patients With Celiac Disease: A Systematic Review and Meta-Analysis. Am. J. Gastroenterol. 110, 1539–1548. doi:10.1038/ajg.2015.296.

Tian N., Faller L., Leffler D. A., Kelly C. P., Hansen J., Bosch J. A., et al. (2017). Salivary gluten degradation and oral microbial profiles in healthy individuals and celiac disease patients. Appl. Environ. Microbiol. 83. doi:10.1128/AEM.03330-16.

Tjellström B., Stenhammar L., Högberg L., Fälth-Magnusson K., Magnusson K.-E., Midtvedt T., et al. (2007). Gut microflora associated characteristics in first-degree relatives of children with celiac disease. Scand. J. Gastroenterol. 42, 1204–1208. doi:10.1080/00365520701320687.

van de Wal Y., Kooy Y. M., van Veelen P. a, Peña S. a, Mearin L. M., Molberg O., et al. (1998). Small intestinal T cells of celiac disease patients recognize a natural pepsin fragment of gliadin. Proc. Natl. Acad. Sci. U. S. A. 95, 10050–10054. doi:10.1073/pnas.95.17.10050.

Verdu E. F., Galipeau H. J., and Jabri B. (2015). Novel players in coeliac disease pathogenesis: role of the gut microbiota. Nat Rev Gastroenterol Hepatol 12, 497–506. doi:10.1038/nrgastro.2015.90.

Villanacci V., Bassotti G., Liserre B., Lanzini A., Lanzarotto F., and Genta R. M. (2006). *Helicobacter pylori* Infection in Patients with Celiac Disease. Am. J. Gastroenterol. 101, 1880–1885. doi:10.1111/j.1572-0241.2006.00621.x.

Ze X., Duncan S. H., Louis P., and Flint H. J. (2012). Ruminococcus bromii is a keystone species for the degradation of resistant starch in the human colon. ISME J. 6, 1535–1543. doi:10.1038/ismej.2012.4.

Zeng M. Y., Inohara N., and Nuñez G. (2017). Mechanisms of inflammation-driven bacterial dysbiosis in the gut. Mucosal Immunol. 10, 18–26. doi:10.1038/mi.2016.75.

